# Improved surfactant-assisted one-pot sample preparation for robust convenient single cell proteomics

**DOI:** 10.64898/2025.12.26.696642

**Authors:** Reta Birhanu Kitata, Zhangyang Xu, Nadia Bayou, Rui Zhao, William B Christler, Matthew J Gaffrey, Karl K Weitz, Mara Serena Serafini, Elisabetta Molteni, Vladislav A Petyuk, Massimo Cristofanilli, Tao Liu, Carolina Reduzzi, Tujin Shi

**Author notes:** R.B.K and Z.X. contributed equally to this manuscript. **Corresponding authors**: Dr. Tujin Shi, Biological Sciences Division, Pacific Northwest National Laboratory, Richland, WA 99352, **Email:**, Dr. Reta Birhanu Kitata, Biological Sciences Division, Pacific Northwest National Laboratory, Richland, WA 99352, **Email:**.

## Abstract

Recent advances in mass spectrometry-based single-cell proteomics (SCP) technologies have revolutionized the SCP field for comprehensive characterization of cellular heterogeneity. However, most of the current SCP approaches employ sub-µL to 1 µL processing volume for effective single-cell sample preparation using either ultralow-volume specialized devices or a 384-well plate by frequently adding water to the plate well to compensate water evaporation, which greatly limits their broad accessibility and analytical robustness. Here we report a robust convenient SCP method termed iSOP (**i**mproved **S**urfactant-assisted **O**ne-**P**ot processing) for processing of single cells at the low µL processing volume using the 384-well plate with tight sealing to avoid sample drying loss. This iSOP SCP method was built upon our previously developed SOP method by the replacement of a PCR tube or 96-well plate with the low-volume 384-well plate and systematic optimization of the single-cell processing conditions. After optimization, 3 µL was selected as the processing volume with a mixture of 2 ng trypsin and 2 ng Lys-C enzymes in terms of robustness, detection sensitivity, and operation convenience. With a commonly accessible LC-MS platform, iSOP-MS can detect and quantify ∼1,200–1,800 protein groups from single HeLa or MCF7 cells. Application of iSOP-MS to two neuroblastoma cell lines has demonstrated that iSOP-MS enabled reliable identification of an average of ∼1,700 and ∼2,050 protein groups from single BE2-C and SK-N-SH cells, respectively, and precise characterization of cellular heterogeneity between the two distinct cell types and within the same cell type. When compared to other available SCP methods, iSOP-MS is more robust and convenient for routine, cost-effective quantitative SCP analysis.

## Introduction

Mass spectrometry (MS)-based proteomics has emerged as a powerful tool for genome-scale proteome profiling.^1–3^ In recent 7 years, there are significant advances in single-cell sample preparation and MS instrumentation for MS-based single-cell proteomics (MS-SCP).^4–12^ With the advanced MS platforms and informatics tools, MS-SCP methods allow for reliable identification of ∼1,500-2,500 proteins from single cells.^4, 9, 13, 14^ With further pushing MS detection sensitivity, the state-of-the-art Orbitrap Astral MS allows identification of ∼5,000 proteins from single cells.^15, 16^ For single-cell sample preparation, specific devices^6, 7, 12, 17, 18^ were designed by greatly downscaling single-cell processing volume down to ∼2-300 nL to minimize sample loss from surface adsorption, but they cannot be easily accessed by research community due to the need of custom-fabricated chips (or capillaries) or a specialized equipment. By contrast, we and others developed multiple convenient methods for cost-effective, easily accessible processing of single cells using standard multi-well plates, PCR tubes, or LC vial inserts to replace the specific devices.^11, 14, 19, 20^ It was demonstrated that they have similar performance as or even better performance than specific device-based methods in terms of technical reproducibility and single-cell proteome coverage.

Despite these advances, current MS-SCP methods still lack robustness and convenience, and there are no standardized protocols for high-throughput SCP analysis. Cell collection and processing with a standard 384-well plate is highly promising for high-throughput SCP. Several 384-well plate-based MS-SCP methods were recently developed with the sub-µL to 1 µL processing volume and slightly different processing procedures (e.g., sealing film or mat and heating temperature).^4, 9, 13–15, 21^ To avoid sample drying for significant loss during single-cell processing, frequent addition of water to the plate wells (e.g., adding 500 nL of water every 15 min) was performed to keep the processing volume at the same level because of a high evaporation rate for sub-µL to 1 µL of water solution in a total of 45 µL plate well.^9, 21^ To further reduce sample drying, single-cell processing in the 384-well plate was operated in a humidity chamber with relatively low protein denaturation temperature and short digestion time.^9, 15, 21^ In addition, single cells may not be sorted in the middle of the well bottom and sub-µL of buffer cannot cover the sorted single cells for effective single-cell processing. Therefore, further improvement of 384-well plate-based MS-SCP methods is needed for robust, convenient, large-scale SCP.

As an early research group pursuing easily accessible MS-SCP methods, in 2021 we developed a cost-effective, broadly adoptable method named SOP-MS, **S**urfactant-assisted **O**ne-**P**ot (SOP) coupled with MS for label-free SCP.^11^ SOP-MS capitalizes on the combination of an MS-compatible nonionic surfactant, n-dodecyl-β-D-maltoside, and a hydrophobic surface-based low-bind PCR tube or 96-well plate for ‘all-in-one’ one-pot sample preparation at the low processing volume of ∼5 μL. This ‘all-in-one’ method, including the elimination of all sample transfer steps and direct injection of single-cell digests from the PCR tube or the 96-well plate, maximally reduces surface adsorption losses for effective single-cell processing. By leveraging our experience in SOP-MS, we have extended SOP-MS to the 384-well PCR plate with systematic optimization of processing volume and enzyme amount to develop an **i**mproved SOP-MS method termed iSOP-MS for robust convenient SCP. iSOP-MS is different from current 384-well plate-based MS-SCP methods without the need of frequent hydration, and it can be operated at normal sample processing conditions. With a commonly accessible Exploris 480 MS platform, iSOP-MS allows reliable identification ∼1,200–2,000 protein groups from single mammalian cells, suggesting comparable performance as other MS-SCP methods.^4, 9, 13, 14^ Application of iSOP-MS to two neuroblastoma cell lines has demonstrated that iSOP-MS enabled precise characterization of cellular heterogeneity between the two distinct cell types and within the same cell type.

### Experimental Section

#### Materials and reagents

Triethylammonium bicarbonate (TEAB), trifluoroacetic acid (TFA), formic acid (FA), acetonitrile (ACN), and n-Dodecyl β-D-maltoside (DDM) were obtained from Sigma-Aldrich (St. Louis, MO). MS-grade water was obtained from Sigma (St. Louis, MO). Promega trypsin gold was purchased from Promega Corporation (Madison, WI), and Lys-C was purchased from FUJIFILM Wako (Richmond, VA). ATCC-formulated Eagle’s minimum essential medium, human recombinant insulin, fetal bovine serum, penicillin/streptomycin, and phosphate-buffered saline (PBS) were purchased from ThermoFisher Scientific (Waltham, MA).

### Cell culture

HeLa, MCF7, BE2-C and SK-N-SH cell lines were obtained from the American Type Culture Collection (ATCC; Manassas, VA). HeLa, BE2-C and SK-N-SH cell lines were cultured and prepared for sorting as described previously.^11, 22^ Briefly, HeLa, BE2-C, and SK-N-SH cells were cultured and maintained in 15 cm Petri dishes in ATCC-formulated Eagle’s minimum essential medium while MCF7 cells were maintained in T-75 flasks and cultured in DMEM/F-12 medium (Corning). The media were supplemented with 0.01 mg/mL human recombinant insulin and a final concentration of 10% fetal bovine serum with 1% penicillin/streptomycin. Cells were maintained at 37 °C in a humidified incubator with 95% O_2_ and 5% CO_2._ Cells were grown until near confluence prior to downstream applications.

### 384-well plate washing and coating

The low-bind 384-well plate was first washed using 20 µL of 0.1% FA in water dispensed into each well using a Mantis liquid handler (Formulatrix), followed by vortexing and complete removal of the liquid. The same washing procedure was repeated using 20 µL of 50% water/ACN. After that 20 µL of 0.005% DDM in 100 mM TEAB was dispensed into each well for coating the plate surface, and the plate was then sealed with a silicone mat and stored at 4 °C until use. Before using the 384-well plate for cell collection, the residual DDM liquid in the plate was discarded and then dried with nitrogen gas in a fume hood.

### Systematic evaluation of sub-µL to low µL water evaporation

Prior to optimization of 384-well plate-based single-cell processing conditions, we have systematically evaluated water evaporation at sub-µL to low µL volumes in the low-bind 384-well plate under standard proteomic sample preparation conditions (i.e., protein denaturation at 75 °C for 1 h and digestion at 37 °C for 10 h). This evaluation was used to determine single-cell sample processing volume. Diluted food-colored water at different volumes of 0.1 µL, 0.2 µL, 0.3 µL, 0.5 µL, 1 µL, 2 µL, 3 µL, 4 µL, 5 µL and 10 µL was dispensed into the 384 plate wells using the Mantis liquid handler. The plate was sealed with a silicone mat and centrifuged at 2,000 rpm for 5 min at 4 °C. The plate was then incubated in a thermocycler (Bio-Rad C1000) at 75 °C for 1 h with the lid temperature set to 95 °C. After incubation, the plate was cooled to room temperature and centrifuged again at 2,000 rpm for 5 min at 4 °C. The plate was placed on ice, and the remaining volume in each well was visually inspected and measured. For the second workflow evaluation, the liquid was subjected to incubation at 75 °C for 1 h with the lid temperature set to 95 °C, followed by heating at 37 °C for 10 h with the lid temperature set to 45 °C. Following incubation, the plate was centrifuged at 2,000 rpm for 5 min at 4 °C and then placed on ice. The remaining liquid volume in the plate well was inspected or accurately measured and the coefficient variation (CV) was calculated.

### Optimization of iSOP with single-cell level input of HeLa cell lysates

For condition optimization, HeLa cell lysates at 0.25 ng protein amount (single-cell level) were used to systematically evaluate different sample processing conditions (different processing volumes and different enzyme amounts and concentrations). To prepare HeLa cell lysate, cultured HeLa cells were rinsed and harvested in ice-cold PBS and cell pellets were resuspended in ice-cold cell lysis buffer (250 mM HEPES, 8 M urea, 150 mM NaCl, 1% Triton X-100, pH 6.0) at a ratio of ∼3:1 lysis buffer to cell pellet. Cell lysates were centrifuged at 14,000 rpm at 4 °C for 10 min, and the soluble protein fraction was retained. Protein concentrations were determined using the BCA assay (Pierce) and then stored at −80 °C in a freezer until further use.

For the evaluation of different processing volumes, the total processing volume was set to 1 µL, 2 µL, 3 µL, 4 µL, 5 µL and 10 µL, with a fixed amount of 2 ng each for both trypsin and Lys-C enzymes. The TEAB buffer was first dispensed into the plate wells at 0 µL, 1 µL, 2 µL, 3 µL, 4 µL and 9 µL using the Mantis liquid handler. 0.5 µL of HeLa lysate at 0.5 ng/µL in 0.4% DDM was then added to each well. The plate was centrifuged at 2,000 rpm for 5 min at 4 °C and incubated in a thermocycler at 75 °C for 1 h with a lid temperature set to 95 °C. After protein denaturation, the plate was cooled down on ice and centrifuged again at 2,000 rpm for 10 min at 4 °C. 0.5 µL of a trypsin/Lys-C mixture at 4 ng/µL for each enzyme was dispensed into each well, followed by centrifugation at 2,000 rpm for 5 min at 4 °C. Digestion was carried out at 37 °C for 10 h with a lid temperature set to 45 °C. The final centrifugation step was performed at 2,000 rpm for 5 min at 4 °C. To quench enzymatic digestion, 5% FA was added as follows: 0.1 µL, 0.2 µL, 0.3 µL, 0.4 µL, 0.5 µL and 1 µL for the processing volumes of 1 µL, 2 µL, 3 µL, 4 µL, 5 µL and 10 µL, respectively, followed by centrifugation at 2,000 rpm for 5 min. Each well was then adjusted to a final volume of 12 µL using 0.1% FA in water. The plate was then centrifuged at 2,000 rpm for 5 min and sealed with a silicon sealing mat. Samples were either analyzed immediately or stored at −20 °C till further LC–MS analysis.

We next evaluated the constant enzyme concentration at 0.66 ng/µL for both trypsin in different processing volumes of 1 µL, 2 µL, 3 µL, 4 µL, 5 µL and 10 µL. Protein denaturation, digestion, and quenching procedures were carried out in the same way as described above for the fixed enzyme amount experiment. The only modification was to dispense 0.5 µL of a trypsin/Lys-C mixture at different concentrations from 1.32 ng/µL to 13.2 ng/µL for each enzyme to have the same enzyme concentration at 0.66 ng/µL for the processing volumes from 1 µL to 10 µL. After processing the single-cell input samples were either analyzed immediately or stored at −20 °C till further LC-MS analysis.

For evaluation of different enzyme amounts, the total enzyme amounts for both trypsin and Lys-C were set to 0.5 ng, 1 ng, 2 ng, 3 ng, 4 ng and 5 ng per well with a fixed processing volume of 3 µL. 2 µL of 0.1% DDM in TEAB buffer was dispensed into each well using the Mantis liquid handler followed by the addition of 0.5 µL of HeLa lysate at 0.5 ng/µL. The plate was centrifuged at 2,000 rpm for 5 min at 4 °C and incubated in a thermocycler at 75 °C for 1 h with a lid temperature set to 95 °C. After the protein denaturation, the plate was cooled down on ice and centrifuged again at 2,000 rpm for 10 min at 4 °C. 0.5 µL of a trypsin/Lys-C mixture at 1 ng/µL, 2 ng/µL, 4 ng/µL, 6 ng/µL, 8 ng/µL or 10 ng/µL for each enzyme was then dispensed into the corresponding wells to achieve the intended total enzyme amounts. The plate was centrifuged at 2,000 rpm for 5 min at 4 °C and subsequently incubated in a thermocycler for digestion at 37 °C for 10 h, with the lid temperature set to 45 °C. The final centrifugation step was performed at 2,000 rpm for 5 min at 4 °C. Enzymatic digestion was quenched by adding 0.3 µL of 5% FA followed by centrifugation at 2,000 rpm for 5 min. Each well was subsequently adjusted to a final volume of 12 µL by adding 8.7 µL of 0.1% FA followed by centrifugation at 2,000 rpm for 5 min. The plate was sealed with a silicon sealing mat. To boost identification, 10 ng of HeLa lysate was similarly processed for generating spectral library at the 3 µL processing volume with six technical replicates. Samples were either analyzed immediately or stored at −20 °C until further LC–MS analysis.

### Single cells and small number of cells sorted by fluorescence-assisted cell sorting (FACS)

Detailed cell preparation before sorting followed our previous publication.^11^ Briefly, after detachment, cells were passed through a 25-gauge needle three times to prevent aggregation. The cells were then suspended in PBS and centrifuged at 500 g for 5 min. This washing step was repeated five times to remove residual trypsin. After that, the cells were resuspended in PBS, filtered through 35 µm mesh caps, and used directly for sorting. For the cell sorting process, a BD Influx flow cytometer (BD Biosciences, San Jose, CA) was used to deposit cells into the 384-well plate containing a pre-dispensed 2 µL of 0.1% DDM in 100 mM TEAB buffer. Alignment of the full-skirted low-bind 384-well plate was done using fluorescent beads (Spherotech, Lake Forest, IL). For each cell line, once sorting gates were established by checking the side scatter detector, cells were sorted into the 384-well plates using the 1-drop single sort mode. In parallel, several blank wells were prepared without cell sorting. After isolation of the desired number of cells, the plate was immediately centrifuged at 2,000 rpm for 10 min at 4 °C to keep the cells at the bottom of the wells. The plates with the isolated cells were then covered with a silicon sealing mat and stored in a −80 °C freezer until further analysis.

### Single cells picked by CellCelector

MCF7 cells were detached from culture flasks using 0.025% trypsin (Gibco), washed and subsequently resuspended in the corresponding culture medium. A total of 5,000 cells in 200 µL of PBS were transferred to a 35-mm Petri dish and allowed to settle by gravity. After cell settling, 3 mL of PBS was gently added to the dish. Individual MCF7 cells were identified by microscopy and isolated by mechanical micromanipulation using the CellCelector (A.L.S.). CellCelector integrates an inverted fluorescence microscope (CKX41 Olympus) equipped with a high-speed automated stage with autofocus capabilities, a CCD camera system (XM10 Olympus), and an automated robotic arm fitted with a glass micro-capillary for cell aspiration allowing for single-cell manipulation.^23, 24^

Automated scanning was performed using a cross-stage speed of 1% and a 20X magnification in the brightfield channel. Image acquisition and microscopic analysis were carried out using CellCelector™ software version 3.1 (A.L.S.). Only spatially isolated single cells were selected for picking to ensure efficient retrieval of one cell per aspiration event. A 15-µm glass capillary was positioned approximately 30 µm above the dish surface, and single cells were aspirated at 1% speed with a volume of 0.1 µL PBS. Aspirated single cells were deposited into the previously washed and coated 384-well plates, as described above, containing 2 µL of 0.1% DDM in 100 mM TEAB buffer per well. Imaging in the brightfield channel (20X magnification) was automatically performed before and after single cell picking to control the collection process and ensure that the targeted cell was successfully retrieved. After isolation of the desired number of cells (n = 30), the plate was immediately centrifuged at 1000 *g* for 10 min at 4 °C to keep the cells at the bottom of the wells. The plates with the isolated cells were then covered with a tight silicon sealing mat, shipped over dry ice and stored in a −80 °C freezer until further analysis.

### Processing of single cells and small numbers of cells in a 384-well plate

The 384-well plate was incubated in a thermocycler at 75 °C for 1 h for protein denaturation with the lid temperature set to 95 °C. The plate was then cooled down on ice and centrifuged at 2,000 rpm for 10 min at 4 °C. After that 1 µL of a trypsin/Lys-C mixture (2 ng/µL trypsin and 2 ng/µL Lys-C) was added to each well followed by centrifugation at 2,000 rpm for 5 min at 4 °C. Digestion was performed at 37 °C for 10 h with the lid temperature set to 45 °C and after digestion the plate remained at 4 °C (idle mode). The plate was then centrifuged at 2,000 rpm for 5 min at 4 °C. Enzymatic digestion was quenched by adding 0.5 µL of 5% FA followed by centrifugation at 2,000 rpm for 5 min. Samples were either analyzed immediately or stored at −20 °C until further LC–MS analysis.

### LC-MS/MS analysis

The Vanquish Neo UHPLC system (ThermoFisher Scientific, San Jose, USA) was coupled with an Orbitrap Exploris 480 MS platform (ThermoFisher Scientific, San Jose, USA). Digested peptides were loaded onto the PepMap Neo Trap Cartridge (ThermoFisher Scientific, San Jose, USA) using Buffer A (0.1% FA in water) at a flow rate of 3 µL/min. The concentrated sample was then separated over a home-packed analytical column (75 µm i.d., 1.7 µm Waters BEH C18, 25-cm Self-Pack PicoFrit column, CoAnn) at a flow rate of 200 nL/min and a 42 min active gradient of 8-40% Buffer B (0.1% FA in acetonitrile). The analysis time is 60 min per sample.

An Orbitrap Exploris 480 MS platform (Thermo Scientific, San Jose, USA) operated at the data-independent acquisition (DIA) mode was used to analyze the separated peptides. Xcalibur software was used to collect MS data. A 2.2 kV high voltage was applied at the ionization source to generate electrospray and ionize peptides. The ion transfer tube was heated to 300 °C to desolvate droplets. High-resolution MS1 data-independent acquisition (HRMS1-DIA) was acquired for proteomic quantification.^13, 25, 26^

For DIA data acquisition, precursor ions from 400 to 760 m/z were scanned at a resolution of 120,000 with a maximum ion injection time set to 125 ms and an AGC target standard. For 1 cell, 5 cells and 10 cells, a fixed precursor isolation window (IW) was set as 24 m/z with 1 Da overlap resulting in 15 scans. Isolated MS1 ions were fragmented using HCD at a 30% level, and MS/MS data were acquired using the following parameters: resolution of 60,000, maximum ion injection time set to auto, AGC target of 1E6, loop control set to 5, and the scan range of 200-1600 m/z. For DIA analysis of 25 cells and 50 cells, MS2 resolution was set to 45,000. For 100 cells DIA data acquisition, a fixed isolation window (IW) was set as 15 m/z with 1 Da overlap with MS/MS resolution of 30,000 and loop control set to 8.

### Data analysis

For DIA data, all the MS raw files were processed in a batch mode and searched together using Spectronaut (Version 19.1) against human reference database (UnipProt 2023-09-01-reviewed with 20,411 entries and 16 contaminants) in default parameters of directDIA+(Deep) workflow with several modifications. Cysteine carbamidomethylation was not included as a modification, while variable modifications were set for methionine oxidation and protein N-terminal acetylation. A trypsin enzyme was selected for cleavage with a maximum number of 2 missed cleavages. Peptide length was set to 7–52 amino acids. Variable modification was set to a maximum of 5, including mass delta 15.9949 for oxidation at M and mass delta 42.0106 for N-terminal acetylation. Precursors were filtered based on 0.01 Q-value cutoff and cross-run normalization was not applied. Quantification was done using MS1 peak area, and filtering for peptides and protein groups was done at 1% FDR cutoff. Downstream data analysis was performed using Perseus (Version 1.6.15).^27^ The non-supervised PCA analysis was used to generate the PCA plot. A Student *t* test was applied to obtain significantly differentially expressed proteins. The extracted data were further processed and visualized using GraphPad Prisim 8 and Microsoft Excel 2017. Signaling pathway enrichment from Reactome database and protein-protein association network were generated using STRING (Version 12.0).^28^

## Results

### iSOP-MS workflow for robust, convenient, label-free single cell proteomics

A high throughput, robust, convenient SCP workflow is required for broad applications of SCP methods for biomedical research. Based on our previous surfactant-assisted one-pot (SOP) processing of single cells and small numbers of cells using a PCR tube or 96-well plate^11, 29, 30^, we further optimized SOP for **i**mproved SOP (iSOP) by processing of single cells in a 384-well plate using a low-cost liquid handler or multi-channel pipette (**Figure 1**). Single cells are directly sorted to the DDM precoated 384-well plate with each well containing 2 µL of preloaded DDM lysis buffer. 3 µL rather than sub-µL volume is used for single-cell processing at standard proteomic sample preparation conditions. The tight sealing of the 384-well plate is created with almost no evaporation for 3 µL by the combined silicone mat and thermocycler. The pre-slit silicone mat allows direct injection of processed single-cell samples without sample transfer. Therefore, 384-well plate-based iSOP-MS can be operated in a robust convenient way for high-throughput SCP analysis.

**Figure 1.**
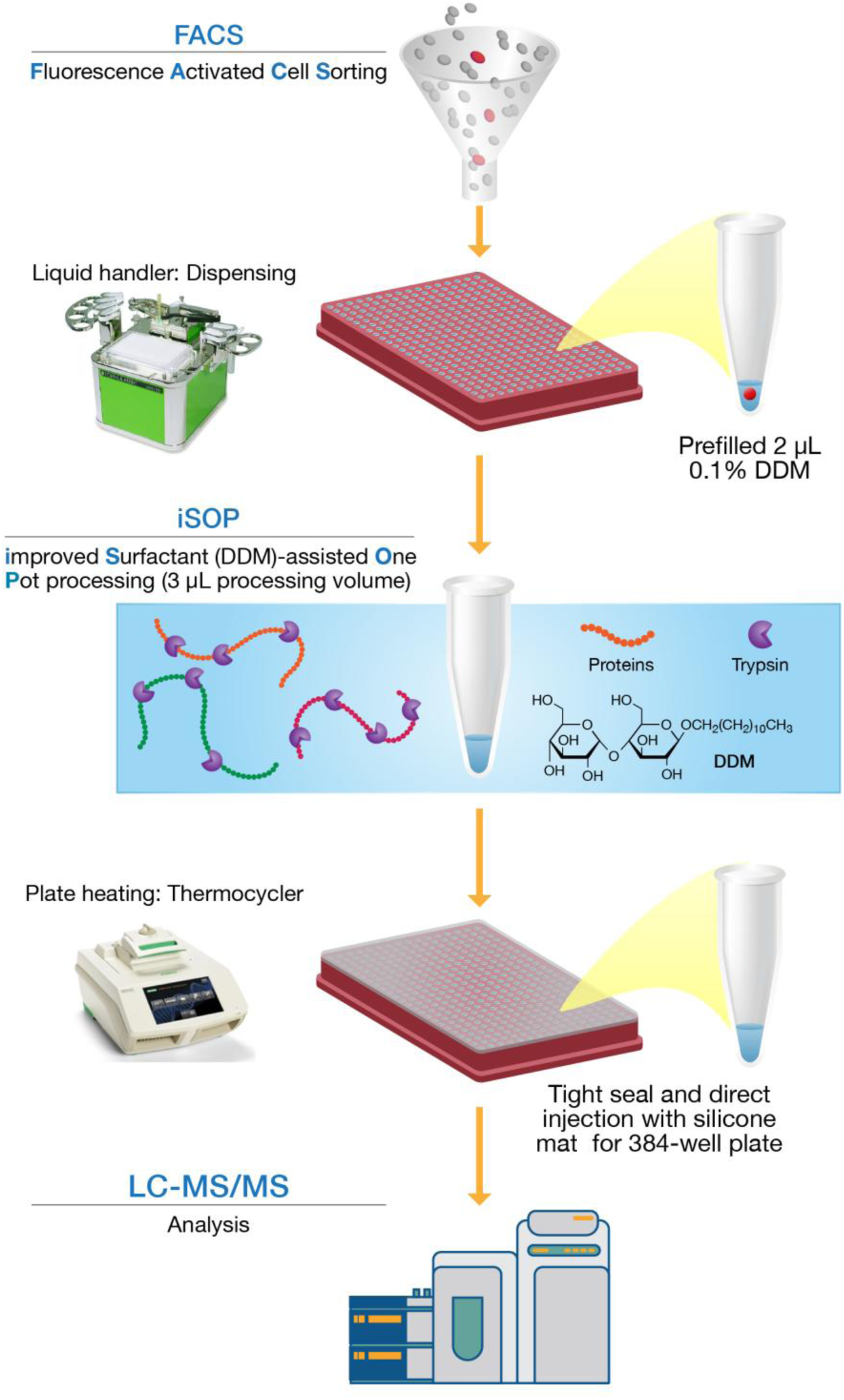
Schematic diagram of the iSOP-MS workflow. Single cells or small numbers of cells are sorted by fluorescence-activated cell sorting (FACS) into a 384-well PCR plate containing preloaded 2 µL of 0.1% (w/v) n-Dodecyl β-D-maltoside (DDM) surfactant in 100 mM TEAB. The sorted cells are subjected to centrifugation at 2,000 rpm for 10 min to ensure that all the sorted cells are at the bottom of the 384-well plate. They are incubated at 75 °C for 1 h over a thermocycler with a silicone mat to cover the plate for protein denaturing. The combination of a silicone mat and thermocycler creates tight sealing with negligible evaporation for the low µL processing volume. After denaturation, 1 µL of a mixture of trypsin and Lys-C at 2 ng/µL per enzyme is dispensed into the 384-well plate and the denatured cells are incubated at 37 °C for 10 h for protein digestion. The peptide digests are directly analyzed by LC–MS/MS for quantitative label-free DIA analysis because the silicone mat used to cover the 384-well plate allows direct automatic injection without sample transfer. A liquid handler (e.g., Mantis) is used for rapid automatic liquid dispensing.

### Systematic evaluation of sub-µL to low µL water evaporation at standard sample processing conditions

Sub-µL to 1 µL processing volume is often used for current 384-well plate-based single-cell proteomics, but it requires frequent hydration and modification of digestion conditions to avoid single-cell sample drying loss. To address this issue, we systematically evaluated water evaporation at standard proteomic sample preparation conditions (protein denaturation at 75 °C for 1 h and digestion at 37 °C for 10 h) for sub-µL to 10 µL (100 nL, 200 nL, 300 nL, 500 nL, 1 µL, 2 µL, 3 µL, 4 µL, 5 µL and 10 µL) commonly used for processing of single cells and small numbers of cells. Two 384-well plates were used to evaluate two conditions with 10 technical replicates per volume and condition: one plate for mimicking protein denaturation and the other plate for mimicking the whole proteomic sample preparation workflow (protein denaturation and digestion). The food-colored water was dispensed into the wells of the 384-well plate using a Mantis liquid handler. The combined silicone mat and thermocycler were used for tight sealing of the 384-well plate. Visual inspecting the wells showed that the bottom center of the wells was covered by the liquid at volumes of 0.5-10 µL while volumes at 100-300 nL only can partially cover the well bottom. For condition 1, after 1 h heating at 75 °C with the lid temperature set to 95 °C, the liquid was completely evaporated for water volumes at 100-300 nL, while ∼0.2 µL, ∼0.8 µL and ∼1.9 µL liquid remained for initial 0.5 µL, 1 µL and 2 µL water, respectively, and the liquid remained almost intact for initial 3 µL to 10 µL water. For condition 2, after 1 h heating at 75 °C (the lid temperature: 95 °C) and then 10 h heating at 37 °C (the lid temperature: 45 °C), the liquid was completely evaporated for water volumes at 100-500 nL, while ∼0.6 µL and ∼1.7 µL liquid remained for initial 1 µL and 2 µL water, respectively, and the liquid remained almost intact for initial 3 µL to 10 µL water. As expected, the CVs were gradually decreased, with large variations for sub-µL volumes, when the water volume increased. Therefore, volumes at 1-10 µL were selected for systematic evaluation of single-cell processing volume and digestion efficiency because they can be operated robustly and conveniently without additional manipulation steps and modification of single-cell processing conditions.

### Systematic evaluation of single-cell processing volume at the fixed enzyme amount or concentration

Based on the above water evaporation evaluation, we have systematically evaluated different processing volumes at 1-10 µL for single-cell level input (0.25 ng of HeLa cell lysate). Our previously established SOP workflow was implemented for this evaluation. For the fixed enzyme amount (a mixture of 2 ng trypsin and 2 ng Lys-C), 1,745 to 2,429 protein groups and 8,463 to 14,119 precursors were detected and identified (**Figure 2A)**. Surprisingly, lower identification numbers for both protein groups and precursors were observed for the processing volumes at 1 µL and 2 µL, which may be attributed to sample loss from evaporation. As expected, with the processing volume increased, the variations in proteome coverage and quantitation reproducibility were gradually decreased and the percentage of the missed cleavage was increased (**Figure 1A**). The 3 µL processing volume resulted in slightly highest precursors (14,119 ± 2,123) and protein groups (2429 ± 277) with a moderate missed cleavage of 10.9 %. Interestingly, the processing volumes from 3 µL to 10 µL have little effect on proteome coverage and quantitation precision for single-cell level sample processing.

**Figure 2.**
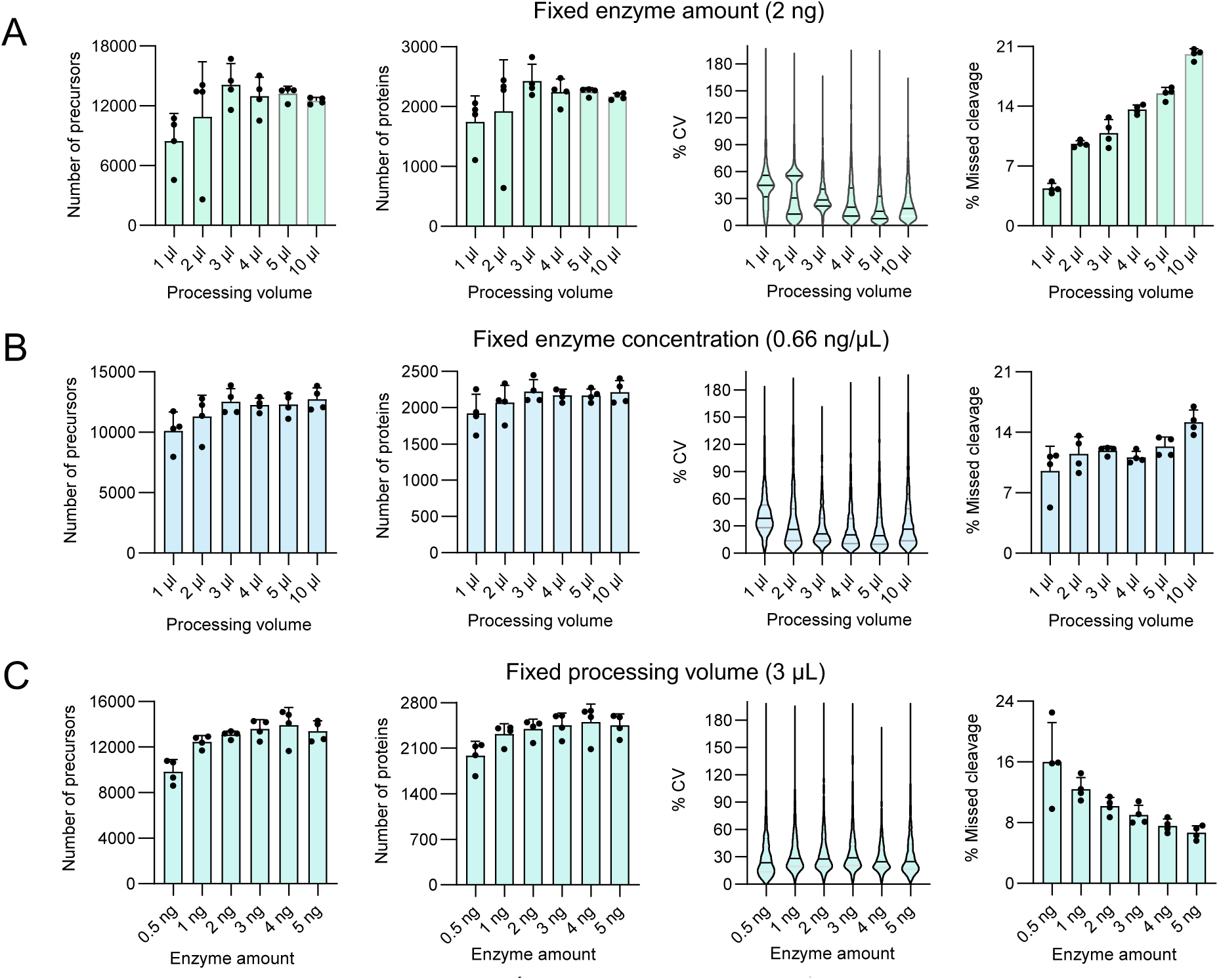
Optimization of iSOP-MS performance using 0.25 ng HeLa cell lysates (equivalent to the protein amount of single cells). (A) Different processing volumes at the fixed enzyme amount (a mixture of 2 ng trypsin and 2 ng Lys-C). (B) Different processing volumes at the fixed enzyme concentration (a mixture of 0.66 ng/µL trypsin and 0.66 ng/µL Lys-C). (C) Different enzyme amounts from 0.5 to 5 ng at the fixed processing volume of 3 µL. The number of peptide precursors and proteins, %CV and % missed cleavage were shown for each condition (Mean ± SD, n = 4 technical replicates).

We have further systematically evaluated different processing volumes at 1-10 µL but with the fixed enzyme concentration (0.66 ng/µL for both trypsin and Lys-C) for 0.25 ng of HeLa cell lysate. Similar as the above evaluation with the fixed enzyme amount, lower identification numbers for both protein groups and precursors were observed for the processing volumes at 1 µL and 2 µL with larger technical variations (**Figure 2B**). For the processing volumes at 3-10 µL, the similar numbers of protein groups and precursors were observed with high technical reproducibility, suggesting that the processing volume has less effect on the proteome coverage and protein quantitation when the surface adsorption is significantly reduced (**Figure 2B**). At the fixed enzyme concentration, the percentage of the missed cleavage is similar with 9.5 to 12.4% for the processing volumes from 1 µL to 5 µL, while the 10 µL processing volume has a relatively higher missed cleavage (∼15%) probably due to sample dilution (**Figure 2B**). All these results suggest that with greatly reduced surface adsorption loss, single-cell level samples can be effectively processed at low µL volumes. Overall, the 3 µL processing volume at the fixed enzyme amount or concentration provided slightly higher number of identified protein groups and precursors with a median missed cleavage (**Figure 2A-B**).

### Systematic evaluation of single-cell digestion efficiency at various enzyme concentrations

With the determined processing volume of 3 µL, next we have systematically evaluated single-cell digestion efficiency at different enzyme concentrations from 0.17 ng/µL to 1.66 ng/µL (i.e., 0.5-5 ng) for each of trypsin and Lys-C enzymes using 0.25 ng of HeLa cell lysate. Except for the lowest enzyme amount (0.5 ng), small difference was observed for the number of identified precursors and protein groups (**Figure 2C**). As expected, the quantitation reproducibility is similar for different enzyme concentrations, and the percentage of the missed cleavage decreased linearly with increased enzyme concentrations (**Figure 2C**). The use of higher enzyme concentrations may interfere with MS signal due to high abundance enzyme peptides from self-digestion, resulting in slightly lower numbers of identified precursors and protein groups at 5 ng of enzyme amount. For bulk sample analysis, a ratio of 1:50 for enzyme:protein is widely used to keep the enzyme concentration at ∼1 ng/µL. There is no systematic evaluation for the enzyme amount in MS-based SCP, but our result showed that ∼1 ng/µL enzyme (i.e., 2 ng trypsin and 2 ng Lys-C in the 3 µL processing volume) would be sufficient to have high digestion efficiency with the percentage of the missed cleavage of <10% and similar proteome coverage as other high enzyme concentrations. With all these systematic evaluations, the 3 µL processing volume with 2 ng for each of trypsin and Lys-C enzymes was finally selected to process sorted single cells.

### iSOP-MS for label-free proteome profiling of single HeLa cells sorted by FACS

After systematic evaluation of single-cell level sample processing conditions, the performance of the optimized iSOP-MS method was evaluated by analysis of single HeLa cells and small numbers of HeLa cells sorted by FACS. 1, 5, 10, 25, 50 and 100 HeLa cells (4 biological replicates per cell number) were sorted directly into a precoated 384-well plate with preloaded 2 µL of 0.1% DDM in 100 mM TEAB. The numbers of identified protein groups were 1,324 ± 242, 2,918 ± 374, 3,112 ± 622, 3,585 ± 71, 4,037 ± 229 and 4,250 ± 28 for 1, 5, 10, 25, 50 and 100 HeLa cells, respectively (**Figure 3A**). High reproducibility in both protein groups and precursors was observed for each cell number (**Figure 3A-B**) and the average protein abundance for commonly quantified 1,834 proteins against the cell number has shown good linearity with R^2^ of 0.8 (**Figure 3C**). Comparative analysis of protein quantitation dynamic range from different cell numbers with the 100 cells have shown that the proteome from single cells is shifted to high abundance proteins, while there is small shift for higher numbers of cells (**Figure 3D**). Furthermore, with similar LC-MS performance, the proteome coverage of single cells from iSOP-MS is comparable to that from other MS-SCP methods.

**Figure 3.**
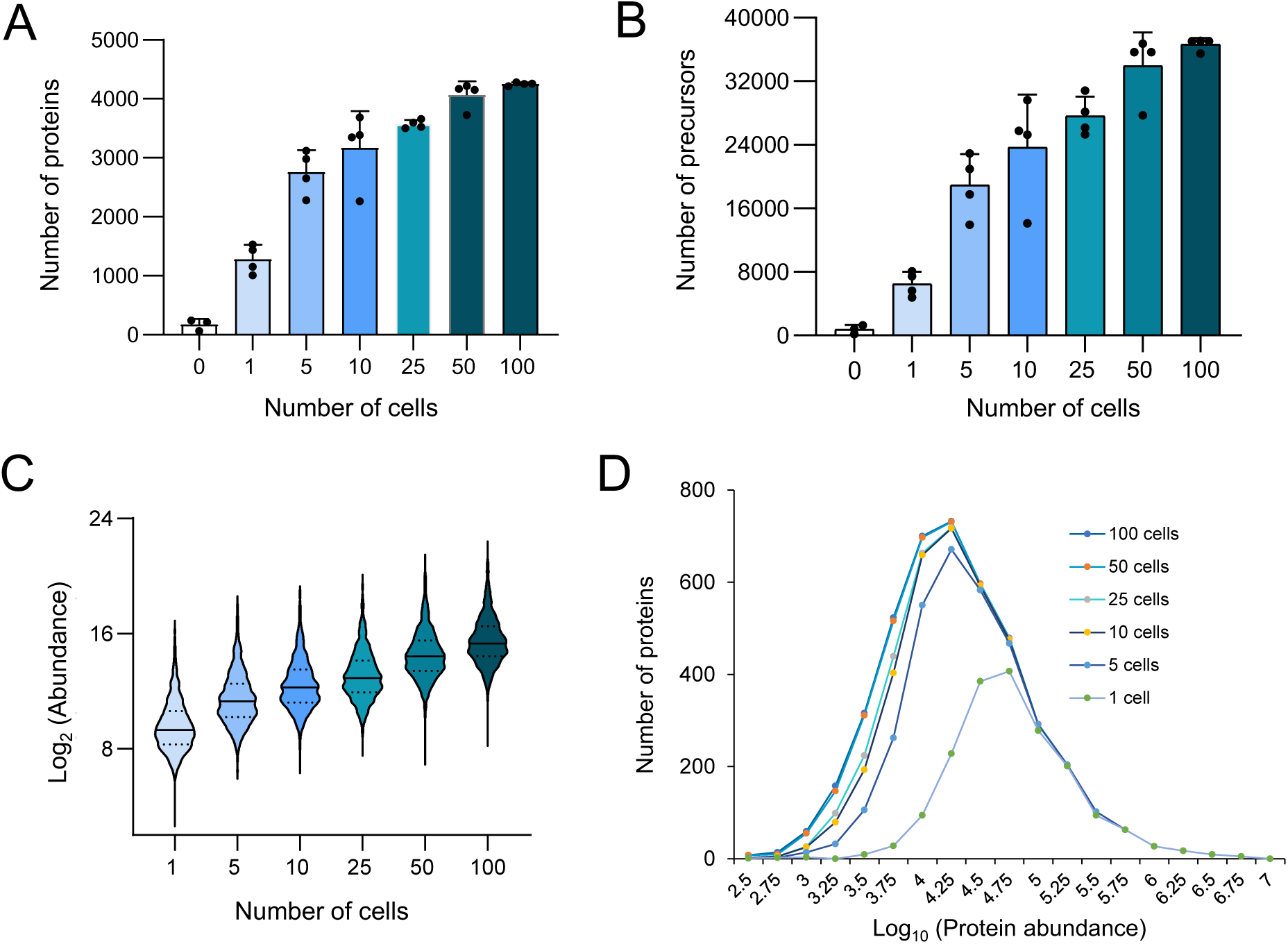
Evaluation of iSOP performance using HeLa cells sorted by FACS. (A) The number of identified protein groups for blank control and 1–100 cells (Mean ± SD, n = 4 technical replicates). (B) The number of identified peptide precursors from blank control and 1–100 cells (Mean ± SD, n = 4 biological replicates). (C) Violin plots showing the distribution of protein abundance at the different number of HeLa cells. Median value is shown by a solid line while the dashed lines show quartiles. (D) The dynamic range of quantified proteins compared to the 100 HeLa cells. Commonly quantified 1,873, 3,421, 3,905, 4,010, 4,296 and 4,313 proteins for 1, 5, 10, 25, 50 and 100 cells were used to plot, respectively.

### iSOP-MS for label-free proteome profiling of single MCF7 cells picked by CellCelector

We next further evaluated the performance of iSOP-MS as well as its transferability for analysis of single MCF7 cells picked by capillary using CellCelector. The CellCelector platform is a fully automated cell imaging and picking system developed for detection, selection and isolation of single cells and clusters of cells (e.g., circulating tumor cells (CTCs)) with high precision.^23, 24, 31^. Single cells are mechanically picked up using the CellCelector and collected in the 384-well plate with preloaded 2 µL of 0.1% DDM in 100 mM TEAB in one laboratory and then shipped to another laboratory for iSOP-MS analysis.

We selected MCF7 breast cancer cell line as a model system for future application of iSOP-MS to CTCs.^32^ Single MCF7 cells were selected from microscope image, automatically picked with glass capillary, and then transferred to the wells of the 384-well plate (**Figure 4A**). 29 single MCF7 cells are successfully picked out of the planned 30 single cells, and they were analyzed by iSOP-MS. A total of 2,372 proteins were quantified from 29 single cells with a median of 1,404 proteins. The cell-to-cell heterogeneity was observed in the proteome coverage and the protein abundance (**Figure 4B-C**). But the subcellular localization analysis has shown that there is no bias in protein distribution across subcellular regions for iSOP-MS (**Figure 4D**). Furthermore, the KEGG pathway analysis has indicated several enriched signaling pathways (n = 88) including estrogen signaling pathway (n = 33 proteins), ERBB signaling (n = 15 proteins), EGFR tyrosine kinase inhibitor resistance (n = 13 proteins), breast cancer pathway (n = 15 proteins) and other pathways dysregulated in breast cancer. All these results demonstrated that iSOP-MS can be flexibly coupled with different cell sorting systems with broad applicability and transferability.

**Figure 4.**
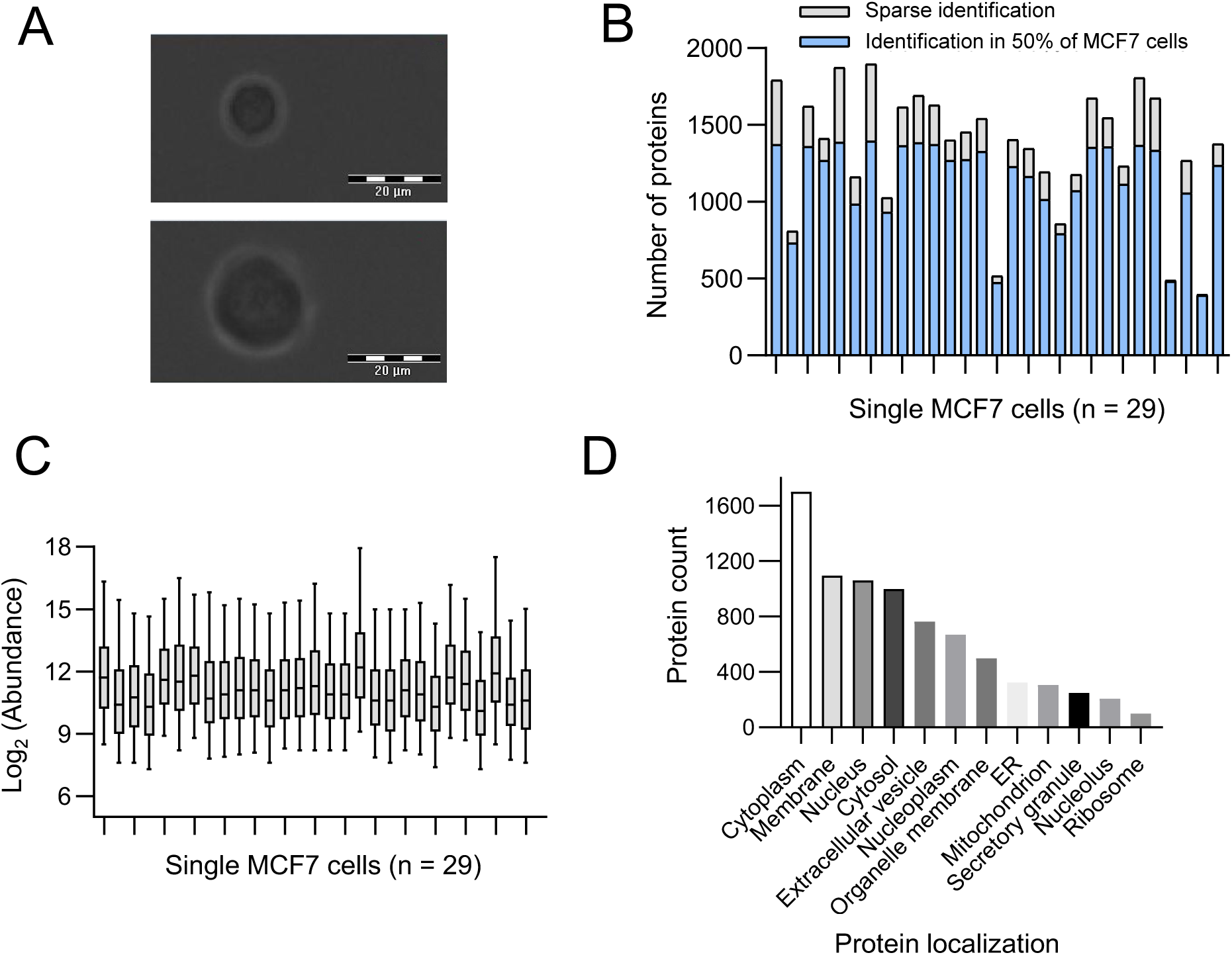
Evaluation of iSOP-MS performance using single MCF7 cells picked by CellCelector (single cell picking by capillary). (A) Images of sorted single MCF7 cells (small and large cells). (B) The number of identified protein groups from single MCF7 cells (n = 29). (C) Protein abundance distribution for each MCF7 cell (boxplot showing 5-95% quartile). (D) Subcellular localization of identified protein groups from gene ontology using DAVID. Single MCF7 cells were sorted into the precoated 384-well plate containing 2 µL of 0.1% DDM (one cell per well) using CellCellector (a microscope-based micro-capillary sorting system).

### Application of iSOP-MS for analysis of single cells from human neuroblastoma cell lines

Neuroblastoma (NB) is the most common pediatric tumor of the developing sympathetic nervous system responsible for ∼8% of all childhood cancer cases with a hallmark of high clinical heterogeneity.^33, 34^ For a proof-of-concept application demonstration, we applied iSOP-MS to characterize two distinct NB cells at single cells and 100 cells, BE2-C and SK-N-SH. The SK-N-SH cell line was found to display epithelial morphology, while the BE2-C cell line is a more aggressive and stem-like phenotype with potential differences in proliferation rate, tumorigenicity, and response to treatment. iSOP-MS allows to identify 1,655 ± 427 and 2,035 ± 683 protein groups from single BE2-C and SK-N-SH cells with 4 biological replicates, respectively (**Figure 5A**). For 100 cells, 3,647 ± 15 and 3,470 ± 128 protein groups were identified from BE2-C and SK-N-SH cells, respectively (**Figure 5A**). Almost all the proteins identified from single cells were covered by the proteome obtained from 100 cells. As expected, principal component analysis has shown higher heterogeneity within single cells than 100 cells (**Figure 5B**). Differential abundance analysis of the two different cell types revealed 2,088 differentially expressed proteins (DEPs) in 100 cells and 256 proteins in single cells (two sampled t-test, p<0.05) with 70% of DEPs in single cells overlapping with the DEPs in 100 cells (**Figure 5C**). Mapping the upregulated proteins in single cells to Reactome database resulted in enrichment of 12 signaling pathways and most of them are cancer associated pathways.

**Figure 5.**
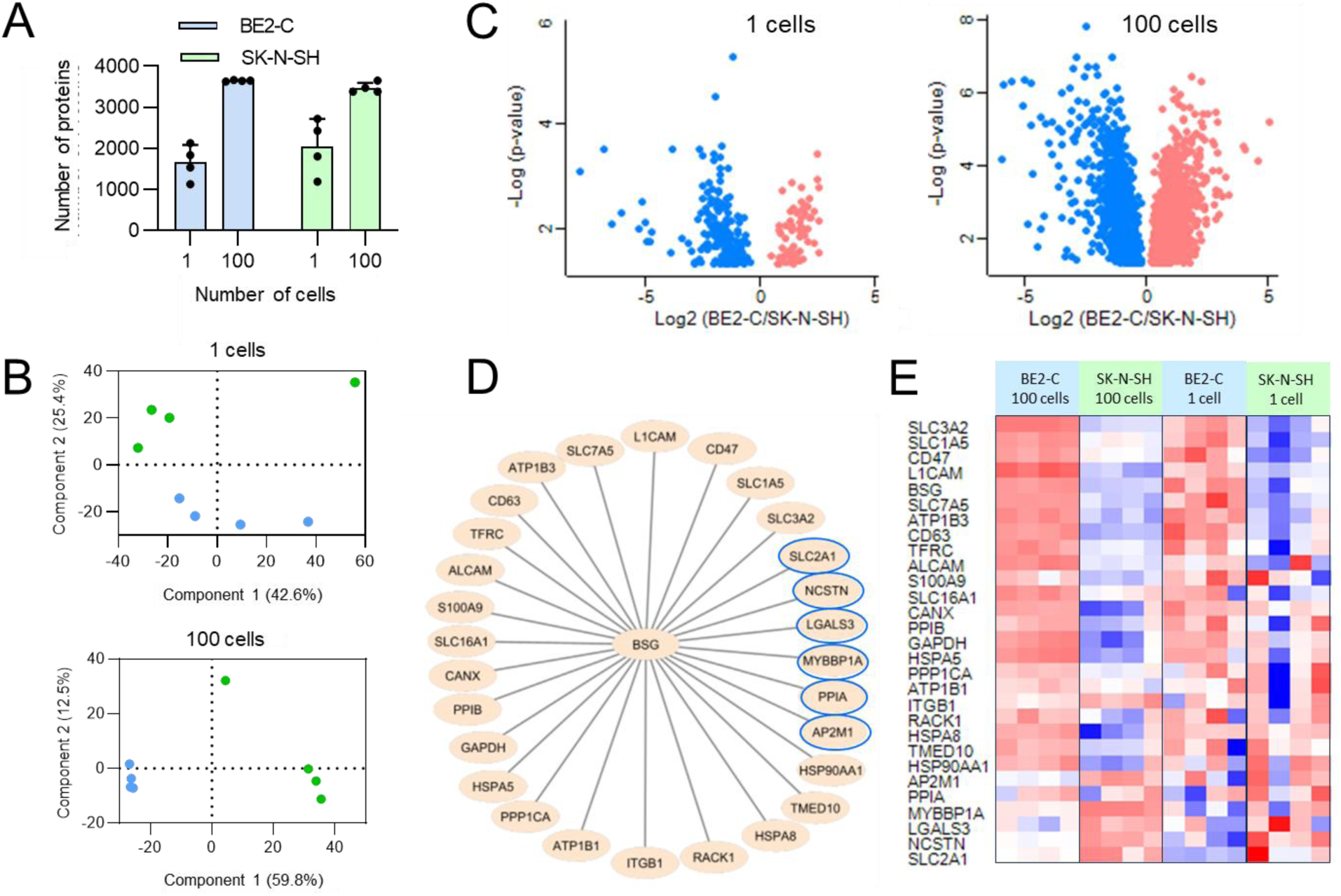
Application of iSOP-MS to FACS-sorted human neuroblastoma cells (BE2-C and SK-N-SH). (A) The number of identified protein groups from single and 100 cells (Mean ± SD, n = 4 replicates). (B) Principal component analysis (PCA) of protein expression levels from single and 100 cells. (C) Volcano plots showing differentially expressed proteins (DEPs) (two sampled t-test, p-value <0.05) between BE2-C and SK-N-SH at single and 100 cells. (D) Protein-protein interaction network of basigin (CD147) signaling based on DEPs. (E) The heatmaps of basigin signaling related proteins showing similar expression trend between single and 100 cells.

CD147, also known as basigin/EMMPRIN (BSG), is a transmembrane protein known to be upregulated in various cancer types^35, 36^ and interacting with several oncogenic proteins to promote solid tumor growth, invasion and metastasis.^36–38^ Protein-protein interaction mapping showed 40 proteins having direct link with CD147 and 29 out of the 40 proteins are upregulated in 100 BE2-C cells compared to SK-N-SH cells (**Figure 5E**). Among the 29 proteins there are 9 proteins observed to be upregulated in single BE2-C cells: BSG (CD147), CD63, CD47, L1CAM, CANX, SLC1A5, SLC3A2, SLC7A5 and ATP1B3 (**Figure 5F**). The interacting partner proteins, SLC3A2, CD63 and L1CAM, were reported to play key role in tumor proliferation and metastasis.^37, 39–41^ Previous analysis of exosomes in NB cell lines also reported higher expression of CD147 and other interacting proteins, consistent with our study.^42^ Immunohistochemistry analysis revealed that the increased CD147 expression was associated with poor overall survival of patients with glioblastoma.^43^ All these results demonstrated the broad utility of iSOP-MS for precise characterization of cellular heterogeneity. Further exploration of the role of CD147 in single NB cells is crucial for providing new insights into NB disease biology.

## Conclusions

We report a 384-well plate-based iSOP-MS method for robust convenient SCP analysis. This method was built upon our previously developed SOP-MS method with systematic optimization of sample processing conditions and the tight sealing of the 384-well plate using the combined 384-well plate silicone mat and a thermocycler heating device. Unlike current 384-well plate-based MS-SCP methods, iSOP-MS can be performed at the low µL processing volume by following a typical proteomic sample processing procedure. With its convenient features, iSOP-MS can be readily implemented for SCP analysis using either a pipette or low-cost liquid handler in any MS laboratory. With further improvements in detection sensitivity and sample throughput, iSOP-MS may open an avenue for routine large-scale SCP analysis with the potential to bridge the gap between single-cell proteomics and transcriptomics technologies.

## Acknowledgements

This work was supported by NIH R01GM139858 (to T.S.), NIH R33CA287139 (to T.S.), NIH RF1MH128885 (to T.S.) and NIH UH3CA256967 (to T.S.). PNNL is a multi-program national laboratory operated by Battelle for the Department of Energy (DOE) under Contract DE-AC05-76RLO 1830. A portion of this research was performed using EMSL, a national scientific user facility sponsored by the DOE’s Office of Biological and Environmental Research and located at PNNL. We thank N.A. Johnson for assistance with graphics.

## Author Contributions

T.S. conceived and designed the study. R.B.K. and T.S. conceptualized iSOP-MS and designed the experiments. R.B.K., Z.X., R.Z. and K.K.W. performed all proteomics experiments and data analysis. N.B., M.S.S., E.M. and C.R. performed MCF7 cell culturing and later MCF7 cell picked by CellCelector. W.B.C. conducted FACS experiments for cell sorting from HeLa and two neuroblastoma cells (BE2-C and SK-N-SH). M.J.G. performed cell culturing for HeLa, BE2-C and SK-N-SH cells. V.A.P. provided two neuroblastoma cell lines. M.C., T.L. and C.R. provided input on the experimental design, data presentation, and manuscript preparation. R.B.K. and T.S. wrote the manuscript with input from all other authors.

